# Integrative analysis of Myrcene’s lipase inhibition and anti-inflammatory mechanisms in human monocytic cells

**DOI:** 10.64898/2026.01.12.699177

**Authors:** Smruti Ranjan Das, Satyabrata Acharya, Sachidanand Nayak, Asenath Shishak, Aditya Ankari

## Abstract

Myrcene (MCN), a monoterpene commonly present in essential oils of aromatic plants, was investigated for its dual role in modulating lipid metabolism and inflammation in human monocytic THP-1 cells. MCN was first evaluated for its ability to inhibit pancreatic lipase activity, followed by cytotoxicity screening to determine non-toxic working concentrations. Its anti-inflammatory effects were assessed by measuring TNF-α gene expression and secretion in LPS-stimulated cells using qRT-PCR and ELISA, while NF-κB p65 nuclear translocation was examined through immunofluorescence. The influence of MCN on macrophage differentiation was further determined in PMA-treated THP-1 cells. MCN significantly reduced pancreatic lipase activity and suppressed inflammatory responses by downregulating TNF-α, TLR2, and MIP-1, alongside preventing NF-κB p65 nuclear migration. Molecular docking supported these findings by confirming favorable interactions of MCN with the catalytic pocket of lipase and the nuclear localization region of NF-κB. Collectively, this study identifies MCN as a promising multitarget natural compound with potential therapeutic application in metabolic and inflammation-associated disorders.

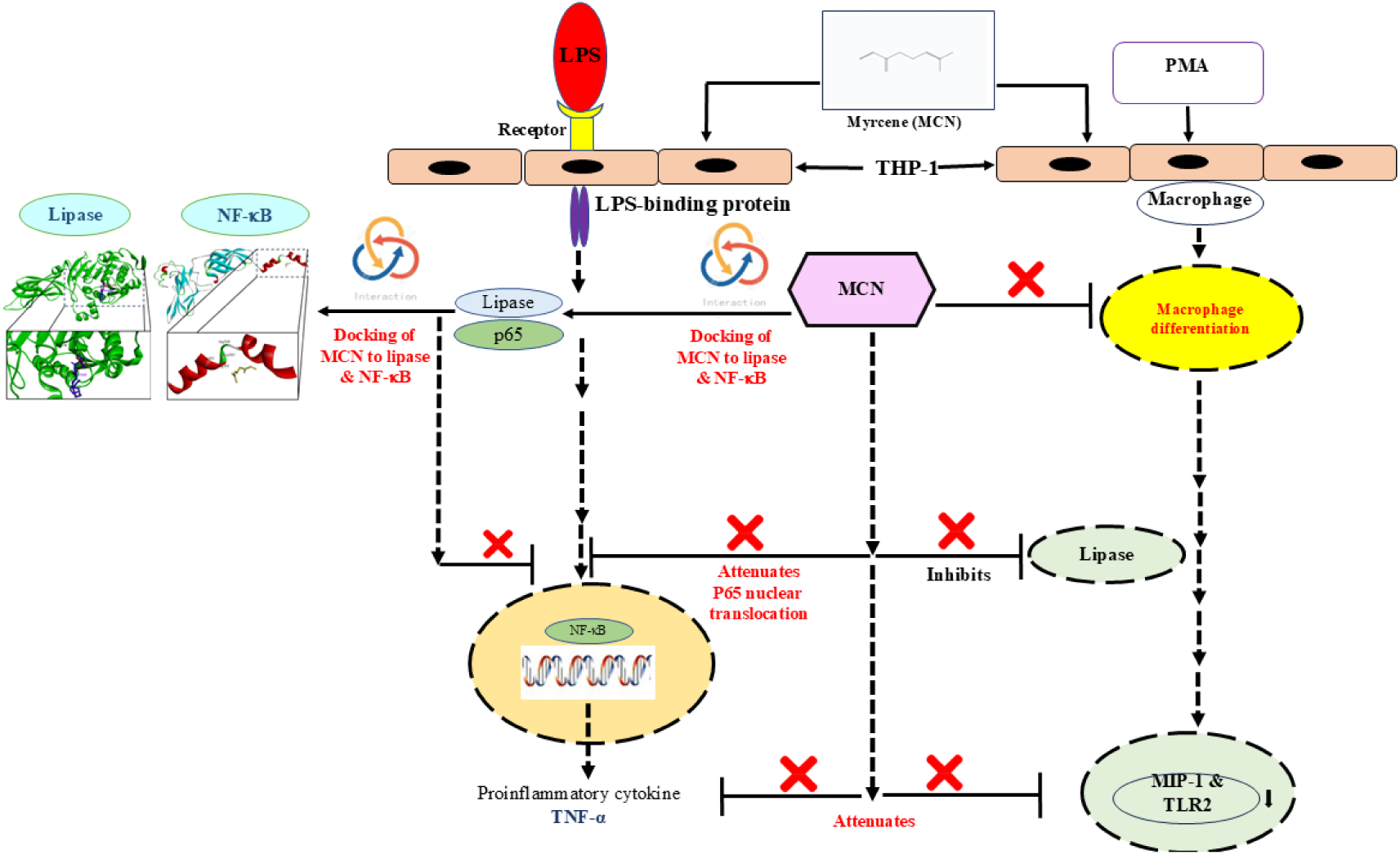

## Introduction

Inflammation is an immunological response of the body against several factors, including dietary lipids, mechanical injury, burns, allergens and noxious stimuli [1]. Usually, it is a defense mechanism of the body; however, chronic inflammation may cause damage to the normal functioning, especially in the case of autoimmune diseases, atherosclerosis, arthritis, etc [2]. Cytokines and chemokines are signaling proteins that play a central role in initiating, amplifying, and resolving inflammation. Pro-inflammatory cytokines such as tumor necrosis factor-α (TNF-α), interleukin-1β (IL-1β), and interleukin-6 (IL-6) stimulate vascular endothelial cells, enhance the expression of adhesion molecules, and recruit immune cells to the site of tissue injury or infection [3]. Chemokines, a specialized subset of cytokines, guide the directed migration of leukocytes by establishing chemotactic gradients. Together, these mediators coordinate the activation of macrophages, neutrophils, and lymphocytes, promoting pathogen clearance and tissue repair [4]. However, excessive or dysregulated production of cytokines and chemokines can result in chronic inflammation, leading to tissue damage and the progression of inflammatory disorders such as rheumatoid arthritis, sepsis, and atherosclerosis [5–7].

Myrcene, a naturally occurring monoterpene widely distributed in essential oils of plants such as lemongrass, hops, thyme, and bay, has been extensively recognized for its diverse pharmacological properties, particularly its strong anti-inflammatory potential. Several experimental investigations, both *in vitro* and *in vivo*, have demonstrated that β-myrcene effectively modulates inflammatory responses by targeting key molecular mediators and signaling pathways. It significantly suppresses the production of major pro-inflammatory cytokines, including tumor necrosis factor-alpha (TNF-α), interleukin-1 beta (IL-1β), and interleukin-6 (IL-6), while concurrently inhibiting the expression of inducible nitric oxide synthase (iNOS) and cyclooxygenase-2 (COX-2), enzymes crucial in the synthesis of inflammatory mediators [8,9].

In murine models of dextran sulfate sodium (DSS)-induced colitis, β-myrcene treatment markedly attenuated intestinal inflammation by reducing disease severity, preserving tissue integrity, and limiting myeloperoxidase activity. Mechanistic studies further revealed that these protective effects were mediated through the inhibition of nuclear factor kappa B (NF-κB) p65 subunit translocation into the nucleus and suppression of mitogen-activated protein kinase (MAPK) signaling cascades, including ERK, JNK, and p38 [8]. In systemic inflammatory conditions, such as diabetic nephropathy and adrenalectomy-induced oxidative stress, myrcene demonstrated significant antioxidant and anti-inflammatory benefits by restoring the activities of key antioxidant enzymes, superoxide dismutase (SOD), catalase (CAT), and glutathione (GSH) and by reducing lipid peroxidation marker malondialdehyde (MDA), thereby protecting tissues from oxidative injury [10,11].

Furthermore, in cell-based studies using human chondrocytes, myrcene mitigated interleukin-1β-induced inflammation by reducing nitric oxide release and downregulating the expression of matrix metalloproteinases MMP-1 and MMP-13, enzymes implicated in cartilage degradation and joint inflammation [12]. These findings collectively suggest that β-myrcene exerts its anti-inflammatory activity through a multifaceted mechanism involving suppression of pro-inflammatory mediators, inhibition of NF-κB and MAPK pathways, and enhancement of the cellular antioxidant defense system. Such pharmacological versatility positions myrcene as a promising bioactive compound for the development of natural anti-inflammatory and cytoprotective therapeutics.

This study aimed to evaluate the inhibitory effect of the compound on pancreatic lipase activity and to assess its anti-inflammatory potential in human monocytic THP-1 cells using *in vitro* assays.

## Materials and methods

### Chemicals

Porcine pancreatic lipase, orlistat, 4-nitrophenyl octanoate, myrcene, DEPC-treated water, Triton X-100, isopropanol, chloroform, Dimethyl sulfoxide (DMSO), Potassium phosphate monobasic, Phorbol 12-myristate 13-acetate (PMA), and trypan blue were purchased from Sigma-Aldrich (Germany). Fetal bovine serum (FBS), Pen Strep and RPMI 1640 medium were purchased from GIBCO (USA). 3-(4,5-dimethylthiazol-2-yl)-2,5-diphenyltetrazolium bromide (MTT), 4’,6-diamidino-2-phenylindole (DAPI), Alexa Fluor 594 goat anti-rabbit IgG (H+L), and Trizol reagent were purchased from Invitrogen (USA). 1X Power SYBR green PCR mastermix was purchased from Applied Biosystems (UK). iScript cDNA synthesis kit was purchased from Biorad (Hercules, CA). NF-κB p65 subunit polyclonal antibody was purchased from Pierce, Thermo Fisher Scientific (USA). Bovine serum albumin (BSA) and p-nitrophenol were purchased from SRL (Mumbai, India). Potassium chloride, di-Potassium hydrogen phosphate, sodium chloride, and sodium nitrate were purchased from Merck (Germany). All other chemicals of analytical grade were purchased from Himedia (Mumbai, India).

### Lipase inhibitory activity

To determine the lipase inhibitory potential of MCN, porcine pancreas lipase enzyme (1mg/1mL in potassium phosphate buffer, pH 7.0) was used as the source. 4-nitrophenyl octanoate was used as the substrate and was solubilized in 100% absolute ethanol. Orlistat was used as a positive control, and it was dissolved in DMSO. The absorbance was measured at a wavelength of 405 nm, at a temperature of 26°C in a microplate reader (Thermo Fisher Scientific) for every 10 minutes up to 1 hour [13,14].

### Cell culture

A human leukemia monocytic (THP-1) cell line was procured from the National Centre for Cell Science (NCCS - Pune, India) and maintained as described previously [5]. Cells were treated with quantified dosages of 20-HE, and cell viability was evaluated by MTT colorimetric assay [7]. THP-1 cells were activated for 3 h by inducing with LPS (0.5 µg/mL). For the monocyte to macrophage differentiation study, cells were stimulated with 25 ng/mL of PMA for 2 days and later images were captured under an inverted microscope.

### Cell viability- MTT assay

The effects of different concentrations (36, 72, 108, and 144 µM) of myrcene (MCN) on THP-1 cells were assessed using 3-(4, 5-dimethylthiazol-2-yl)-2,5-diphenyltetrazolium bromide (MTT) assays, following the methodology described by [5]. THP-1 cells were plated at a density of 5 × 10^5^ cells/mL in 24-well plates, treated with varying concentrations of MCN, and incubated at 37°C overnight. After the incubation period, the cells were harvested, and the cell pellet was washed three times with RPMI medium. Subsequently, 20 µL of MTT solution (5 mg/mL) was added to the cells, and the mixture was incubated for 4 hours. The cells were then pelleted by low-speed centrifugation and incubated for 15 minutes with 100 µL of dimethyl sulfoxide (DMSO) to dissolve the insoluble purple formazan crystals. The absorbance of the MTT formazan was measured at λ 570 nm, using λ 690 nm as a reference wavelength, with a 96-well plate reader (Thermo Fisher Scientific) [15,16].

### Quantification of transcripts of TNF-α, MIP-1, and TLR2

Total RNA was isolated from THP-1 cells following the respective treatments using TRIzol reagent, following the manufacturer’s protocol (Invitrogen). Complementary DNA (cDNA) was synthesized from the extracted RNA using the iScript™ cDNA synthesis kit. For the real-time quantitative PCR (qRT-PCR) analysis, 2 µL of cDNA was used as a template along with 1X FG Power SYBR Green PCR Master Mix (Applied Biosystems). Gene expression levels of inflammatory markers, including TNF-α, MIP-1, and TLR2, were quantified using gene-specific primers (Table 1). The specificity of the amplification was verified by generating a melting curve, in which fluorescence was plotted as a function of temperature [5,7,17]. The relative transcript levels of target genes were normalized to those of the housekeeping gene GAPDH to ensure accurate quantification.

**Table 1:**
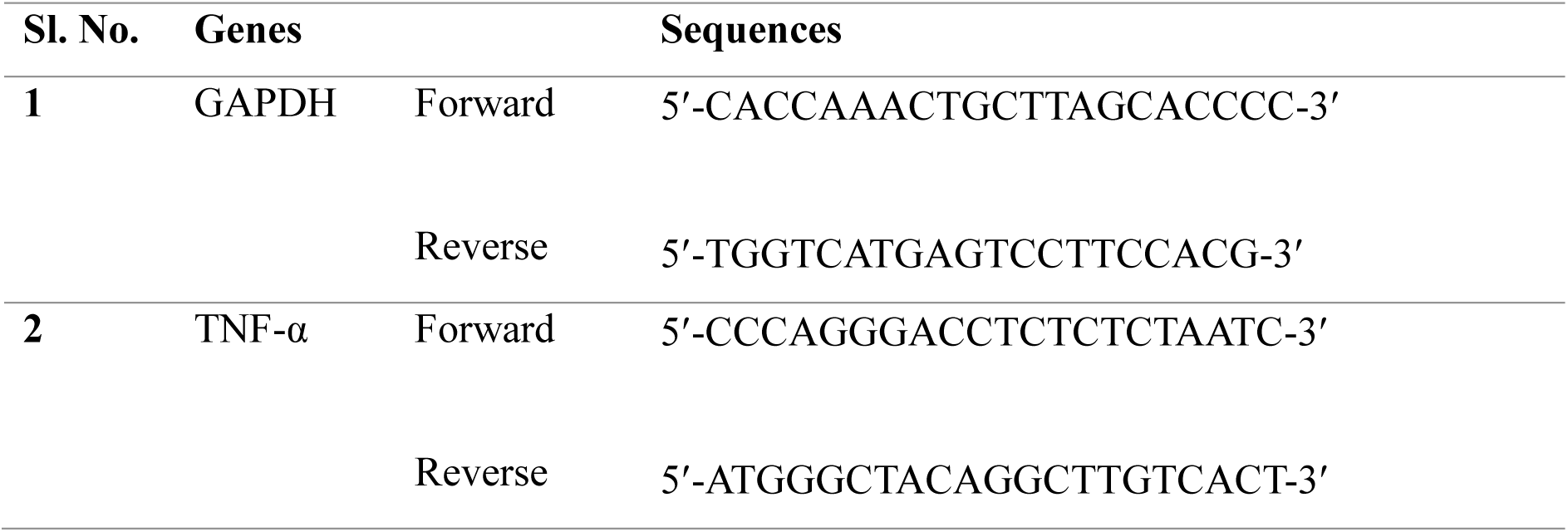
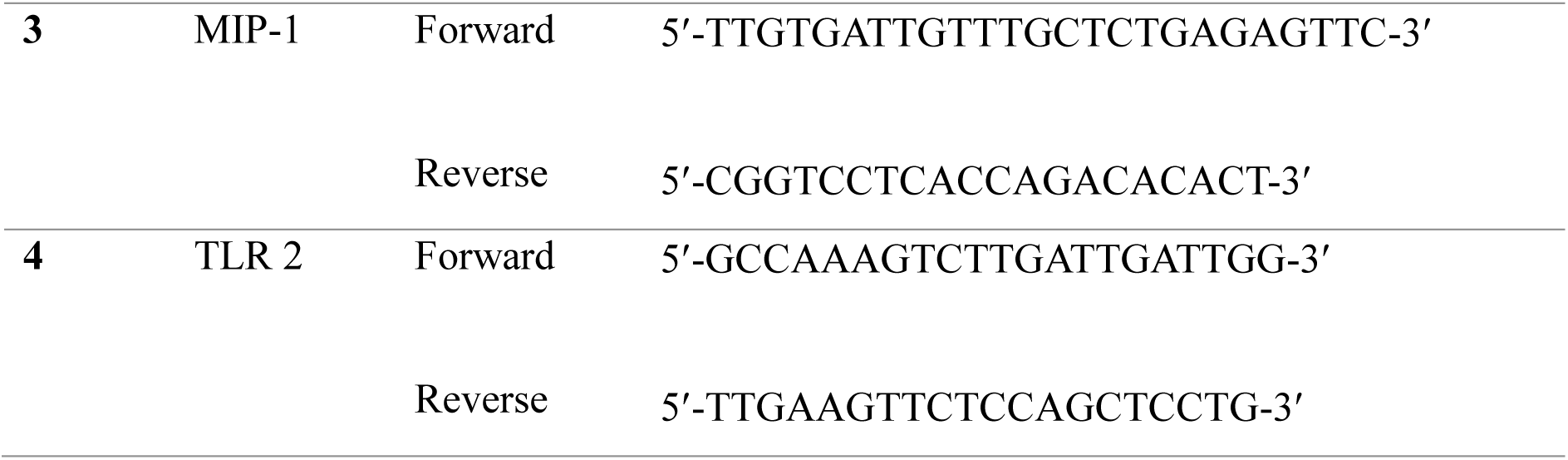
Primers used for real-time quantitative PCR assays.

### Determination of secreted proinflammatory marker TNF-α by the ELISA method

Secreted TNF-α is a proinflammatory cytokine secreted by THP-1 cells, and was measured with BD OptEIA−™ Set Human ELISA kits (San Diego, CA, USA). THP-1 cells at a density of 5 × 10^5^ cells/mL were incubated with and without different concentrations of test compounds overnight and induced with LPS (0.5 µg/mL) on the following day for 3 h [5,7]. After induction, the cells were spun, and the supernatants were collected for quantification of the above marker levels *via* ELISA kits (BD Biosciences). The reaction mixture was assayed by measuring the absorbance at a wavelength of λ 450 nm by a microplate reader with the wavelength correction set at λ 570 nm. Secretory levels in culture supernatants were determined by using a standard curve, which was constructed by serial dilutions of the TNF-α as provided with the ELISA kits.

### Immunofluorescence detection by confocal microscopy

Cells grown on sterile coverslips were fixed with 4% paraformaldehyde for 15 minutes and washed with PBS. Permeabilization was performed using 0.1% Triton X-100 for 10 minutes, followed by blocking with 3% BSA for 1 hour to minimize non-specific binding. Cells were incubated overnight at 4 °C with primary antibodies, washed, and then exposed to fluorescently labelled secondary antibodies for 1 hour at room temperature in the dark. Nuclei were stained with DAPI, and coverslips were mounted using glycerol. Fluorescent images were visualised under a confocal microscope [16].

### PMA (phorbol-12-myristate-13-acetate) stimulated monocyte-derived macrophage differentiation

THP-1 cells grown in serum-free media (2% v/v) were pretreated with MCN and stimulated with PMA (25 ng/mL) for 48 h at 37°C in 5% CO_2_ as described by Dong et al., 2014 [18]. The phenotype of THP-1 cells was observed under an inverted microscope. THP-1 cells without PMA were used as control cells (undifferentiated).

### Molecular docking

The pancreatic lipase-colipase complex (PDB ID: 1LPB) and the NF-κB complex (PDB ID: 1NFI) 3D protein structures were downloaded from the Protein Data Bank (PDB)(https://www.rcsb.org/). All structural components other than NF-κB p65 (Chain A) were deleted from the PDB file (PDB ID: 1NFI) using the PyMOL Molecular Graphics System, Version 3.1.4.1 Schrödinger, LLC and the resulting structure was saved as a new PDB file. Similarly, all structural components other than the lipase (Chain B) were deleted from the PDB file (PDB ID: 1LPB) using the PyMOL Molecular Graphics System, Version 3.1.4.1 Schrödinger, LLC, and were saved as a separate PDB file. Both the resulting PDB files containing NF-κB P65 (Chain A) and lipase (Chain B), respectively, were prepared for molecular docking using the AutoDockTools v1.5.7 [19] separately.

The molecular structure of the myrcene (3D coordinates) (PubChem CID: 31253) and the orlistat (2D coordinates) (PubChem CID: 3034010) were downloaded from the PubChem database (https://pubchem.ncbi.nlm.nih.gov/). Since only the 2D structure/coordinates of orlistat were available in the database, its corresponding 3D structure/coordinates were generated using Open Babel v3.1.1 [20]. Both ligands were prepared for molecular docking using AutoDockTools v1.5.7 [19] separately.

The molecular docking compatible file of orlistat (ligand) and myrcene (ligand) was individually subjected to blind rigid docking with the molecular docking compatible file of lipase (Chain B) (protein) using AutoDock Vina v1.1.2 [21] with ten different random seed values each.

The molecular docking compatible file of myrcene (ligand) was subjected to blind rigid docking with the molecular docking compatible file of NF-κB p65 (Chain A) (protein) using AutoDock Vina v1.1.2 [20] with ten different random seed values.

The molecular docking outputs, including binding affinities and interaction poses, were visualised using the BIOVIA, Dassault Systèmes, Discovery Studio Visualiser, v25.1.0.24284, San Diego: Dassault Systèmes, 2025.

### Statistical analysis

The entire data in the manuscript represents the mean ± standard deviation (SD) of triplicates. Data were analysed by one-way analysis of variance (ANOVA).

## Results

### Lipase inhibitory effect of MCN

Lipase activity reached its maximum within 20 minutes, as evidenced by an increase in absorbance at 405 nm. Under the same experimental conditions, the standard inhibitor orlistat exhibited 33 ± 2% inhibition at a concentration of 100 µg/mL. The inhibitory potential of MCN against pancreatic lipase was then evaluated under identical conditions. MCN demonstrated a concentration-dependent inhibitory effect, with inhibition rates of 14.2 ± 0.06%, 19.5 ± 0.04%, and 43.6 ± 5% at concentrations of 90, 180, and 360 µM, respectively (Fig. 1).

**Fig. 1.**
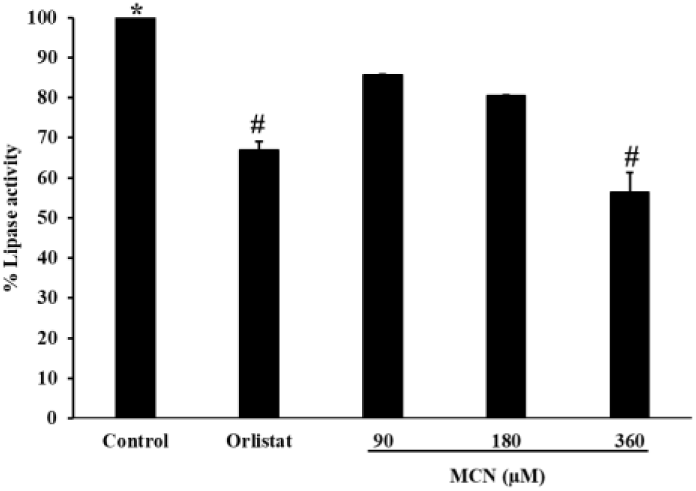
Effect of selected MCN on pancreatic lipase enzyme. Data are presented as mean ± standard deviation (SD) from three independent experiments (n=3); statistical significance was set at P< 0.05.

### Effect of MCN on cell viability of THP-1 cells

The cytotoxicity of the synthesized conjugates on THP-1 cells was determined MTT assay. The THP-1 cells treated with phytocompound MCN at 4 different concentrations (36, 72, 108, and 144 µM) did not have any toxic effects; however, the cell viability was approximately equal to that of the control cells (Fig. 2).

**Fig. 2.**
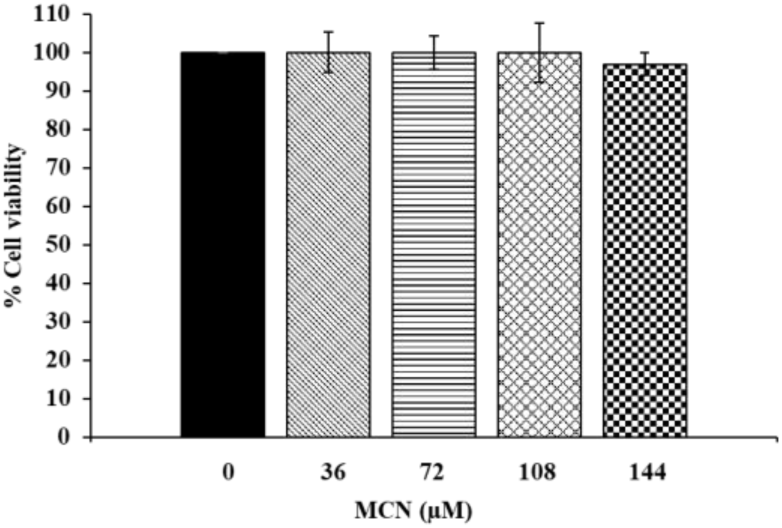
MTT assay of MCN in THP-1 cells. Cell viability was assessed after 15 h of treatment at 37 °C in 5% CO₂ with different concentrations of MCN. Samples with media alone (treated with MCN) were used as blanks. The data are presented as the mean ± SD of 3 independent experiments.

### Effect of MCN on LPS-induced proinflammatory marker in THP-1 cells

The pure metabolite myrcene (MCN) demonstrated a strong inhibitory effect on the expression and secretion of the pro-inflammatory cytokine TNF-α in a concentration-dependent manner. When THP-1 cells were treated with increasing concentrations of MCN (36, 72, 108, and 144 µM), a marked reduction in TNF-α transcript levels was observed, showing 1.5 ± 0.07-fold, 80 ± 2.08-fold, 5.5 ± 0.08-fold, and 3.2 ± 0.5-fold decreases, respectively. In contrast, LPS-stimulated control cells exhibited a substantial upregulation of TNF-α expression by approximately 172 ± 4-fold, confirming an active inflammatory response (Fig. 3a).

**Fig. 3.**
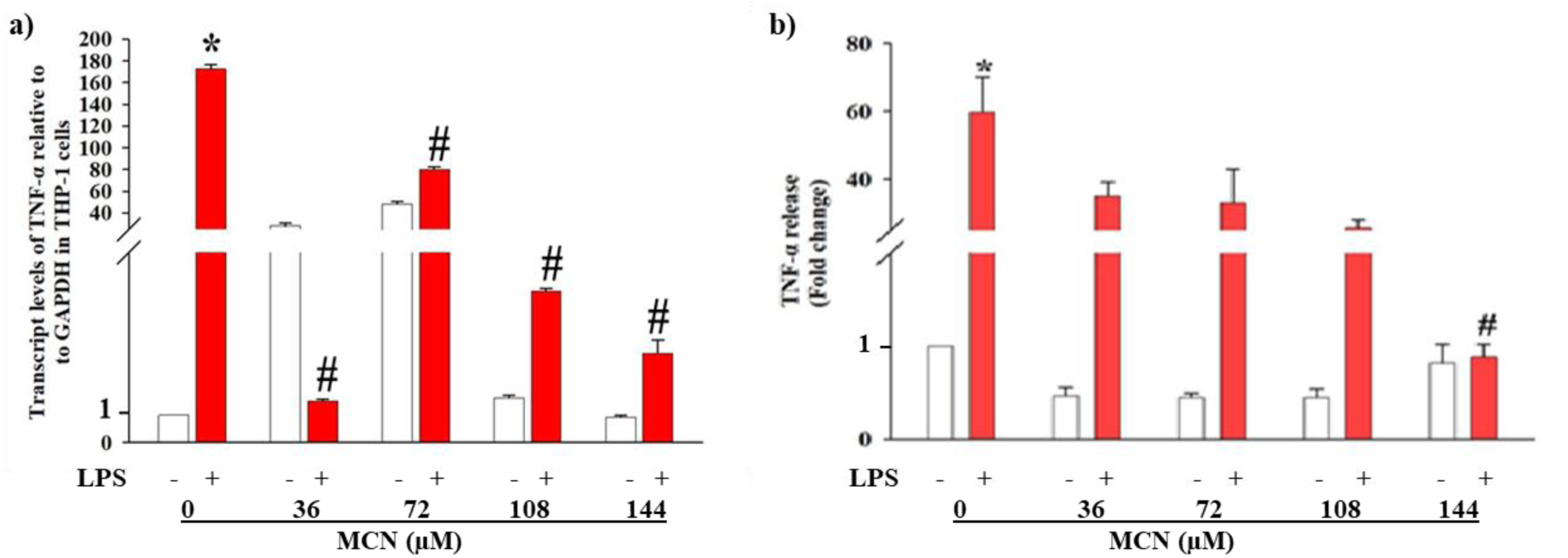
Effect of MCN on LPS-induced TNF-α gene expression and secretion in THP-1 cells. THP-1 cells were pretreated with or without MCN and subsequently stimulated with LPS for 3 hours. Following treatment, (a) TNF-α mRNA expression was analyzed by quantitative real-time PCR, and (b) TNF-α protein secretion in the culture supernatant was measured using ELISA. Data are presented as the mean ± SD from three independent experiments. #p<0.001 indicates a significant difference between MCN-treated and untreated groups, whereas *p<0.001 denotes statistical significance between the control groups.

Furthermore, MCN treatment significantly decreased TNF-α protein secretion in LPS-stimulated THP-1 cells, with the highest concentration (144 µM) reducing TNF-α secretion to 0.88 ± 0.1-fold, whereas the LPS control group showed a 59.8 ± 10-fold increase (Fig. 3b).

These findings collectively indicate that MCN effectively suppresses both the transcriptional and protein levels of TNF-α, thereby mitigating excessive cytokine production. The results suggest that MCN exerts a potent anti-inflammatory effect by modulating LPS-induced signaling pathways and may play a protective role in preventing inflammatory damage in human monocytic cells.

### Effect of MCN on LPS-activated NF-κB nuclear translocation

Based on the observed inhibitory effect of MCN on LPS-induced TNF-α secretion, its influence on NF-κB (p65) nuclear translocation was further investigated in LPS-stimulated human monocytic THP-1 cells. Immunofluorescence analysis revealed that treatment with MCN at concentrations of 108 and 144 µM effectively prevented the nuclear translocation of the NF-κB p65 subunit (Fig. 4b). In LPS-stimulated control cells, the majority of p65 protein was translocated into the nucleus, indicating strong activation of the NF-κB pathway (Fig. 4a). However, in MCN-treated cells, p65 remained predominantly localized in the cytoplasm, suggesting that MCN suppresses NF-κB activation by inhibiting its nuclear translocation and thereby contributes to its anti-inflammatory mechanism.

**Fig. 4.**
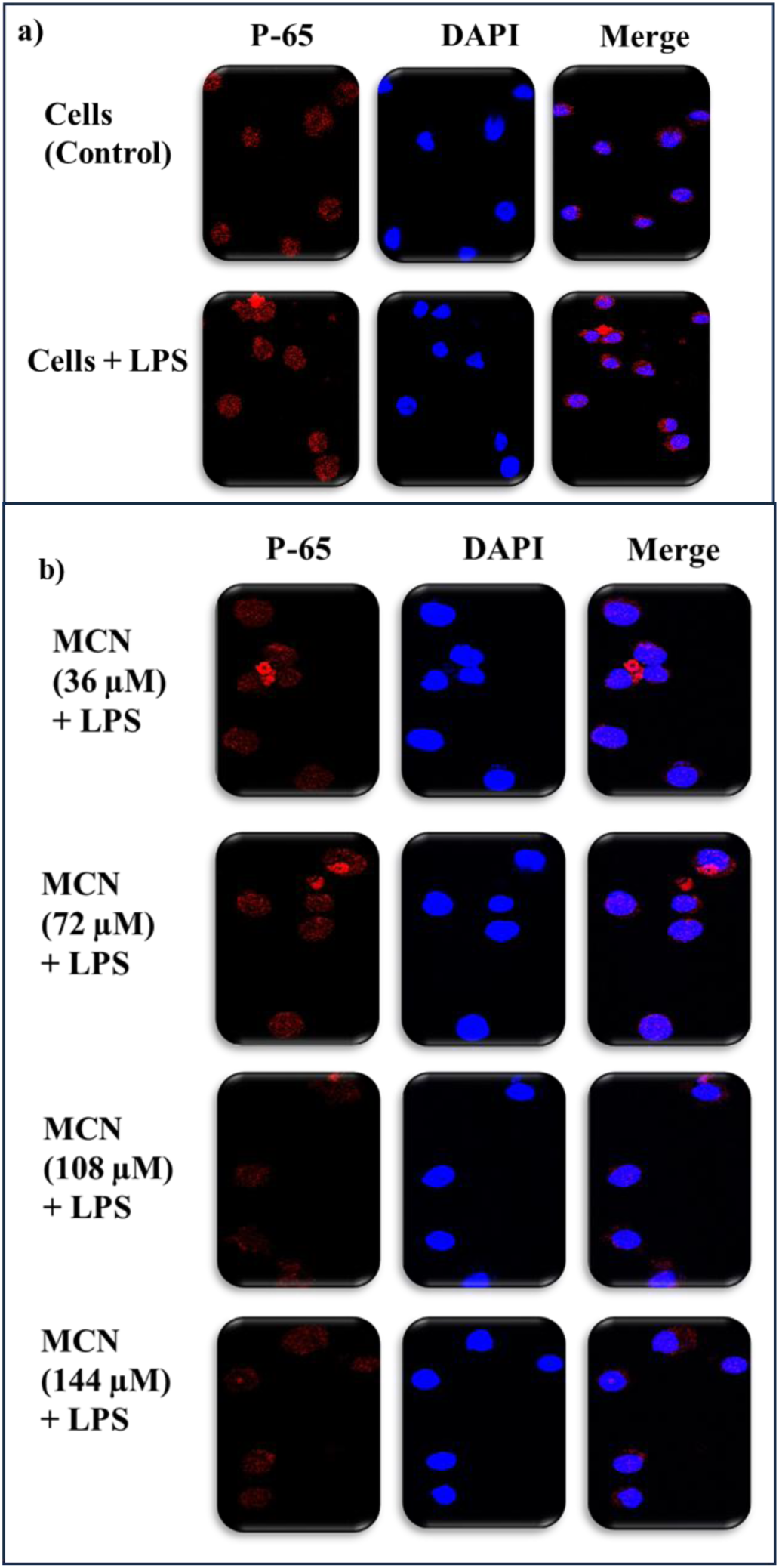
Effect of MCN on LPS-induced NF-κB p65 subunit translocation. THP-1 cells were stimulated with LPS (0.5 µg/mL), and the subcellular localization of the p65 subunit was assessed using immunofluorescence. NF-κB p65 was detected with an Alexa Fluor 594-conjugated secondary antibody, and nuclei were counterstained with DAPI. Images were acquired using a Leica confocal microscope. Panels represent: (a) Control and (b) MCN.

### Effect of MCN on PMA-induced monocyte to macrophage differentiation in THP-1 cells

Human monocytic THP-1 cells are spherical, non-adherent suspension cells that typically do not attach to the substratum of culture plates. Upon stimulation with phorbol 12-myristate 13-acetate (PMA, 25 ng/mL) for 48 hours, these cells undergo differentiation into macrophage-like cells, acquiring an adherent morphology and attaching firmly to the culture plate/dish (Fig. 5a). To assess the effect of MCN on this differentiation process, THP-1 cells were pretreated with varying concentrations of MCN (36-144 µM) for 12 hours, followed by PMA treatment for an additional 48 hours. MCN treatment resulted in a concentration-dependent inhibition of PMA-induced monocyte-to-macrophage differentiation, with a pronounced suppressive effect observed at 72, 108, and 144 µM (Fig. 5b).

**Fig. 5.**
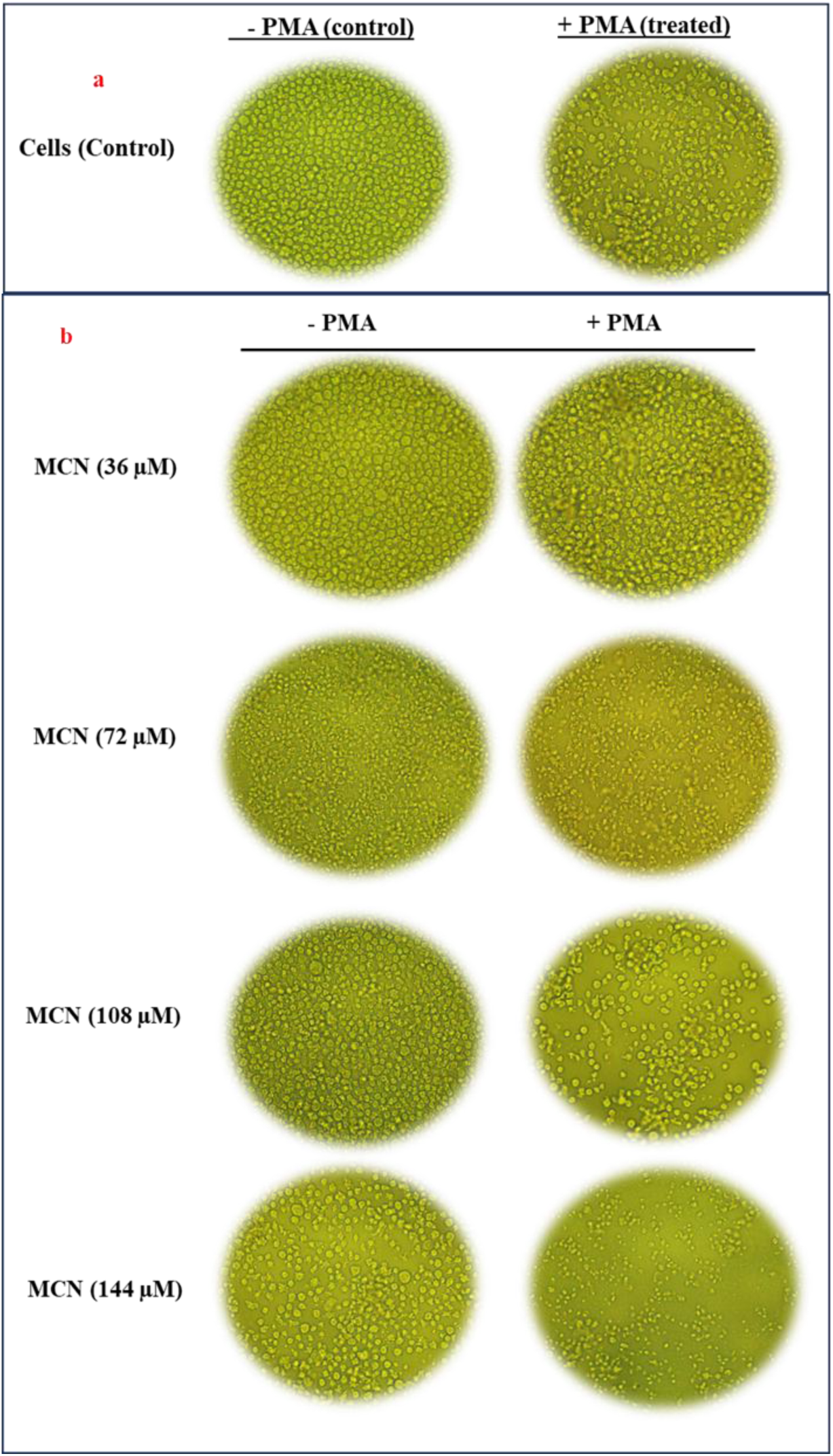
Effect of MCN on PMA-induced monocyte to macrophage differentiation. THP-1 cells were stimulated with MCN at different concentrations, followed by PMA (25 ng/mL) for 48 h. Images were acquired using an inverted microscope (Magnus). Panels represent: (a) Control and (b) MCN.

### Modulation of gene expression during PMA-induced differentiation of THP-1 cells by MCN

Stimulation of THP-1 cells with PMA resulted in a pronounced upregulation of MIP-1 and TLR2 gene expression, showing increases of 17.7 ± 1.2-fold and 2689.04 ± 30.6-fold, respectively, compared to untreated control cells. However, pretreatment with MCN significantly attenuated this PMA-induced response in a concentration-dependent manner. TLR2 expression was markedly suppressed to 76.3 ± 3.1, 276.1 ± 1.2, 261.1 ± 4, and 246.2 ± 6.8 folds at MCN concentrations of 36, 72, 108, and 144 µM, respectively (Fig. 6b). Similarly, MIP-1 expression was reduced to 2.02 ± 0.3 and 2 ± 0.2 folds at 108 and 144 µM, respectively (Fig. 6a). These findings indicate that MCN effectively downregulates PMA-induced inflammatory gene expression, suggesting its potential role in modulating macrophage activation and attenuating the inflammatory response.

**Fig. 6.**
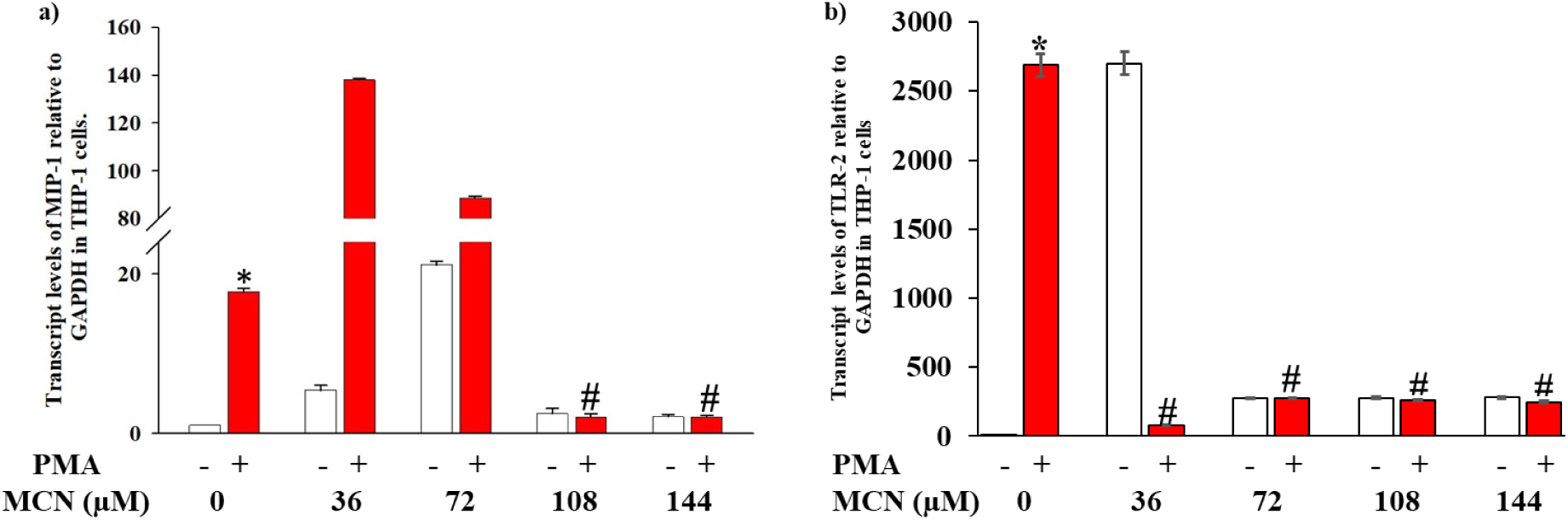
Effects of MCN on PMA-induced MIP-1 and TLR2 gene expression, as determined *via* quantitative real-time PCR. THP-1 cells were pretreated with MCN for 12 h, followed by 48 h of PMA induction. At the end of the treatment, the transcript levels of (a) MIP-1 and (b) TLR2 were quantified *via* real-time PCR. Experiments were performed at least in triplicate, and the results are expressed as the mean ± S.D. *p < 0.001 for comparison between PMA-induced cells and PMA-uninduced cells. #p<0.001 compared between cells treated with PMA in the presence of MCN *vs.* control.

### Molecular docking analysis of pancreatic lipase with MCN

Both myrcene and orlistat (used as the reference inhibitor) were consistently docked within the same active binding pocket of pancreatic lipase (Chain B) across ten independent molecular docking simulations, confirming the reliability of the docking results (Fig. 7a). Orlistat and myrcene exhibited the highest binding affinity of −6.8 kcal/mol and −5.6 kcal/mol, respectively, indicating a favorable interaction of both ligands with the enzyme. As reported in Gholami et al. (2025) [22], the catalytic activity of pancreatic lipase is mediated by a conserved catalytic triad comprising Ser152, Asp176, and His263. In the present analysis, orlistat exhibited direct hydrogen bonding interactions with two residues of this catalytic triad, Ser152 and His263 (Fig. 7a and 7b), while myrcene demonstrated a direct interaction with His263 (Fig. 7a and 7c).

**Fig. 7.**
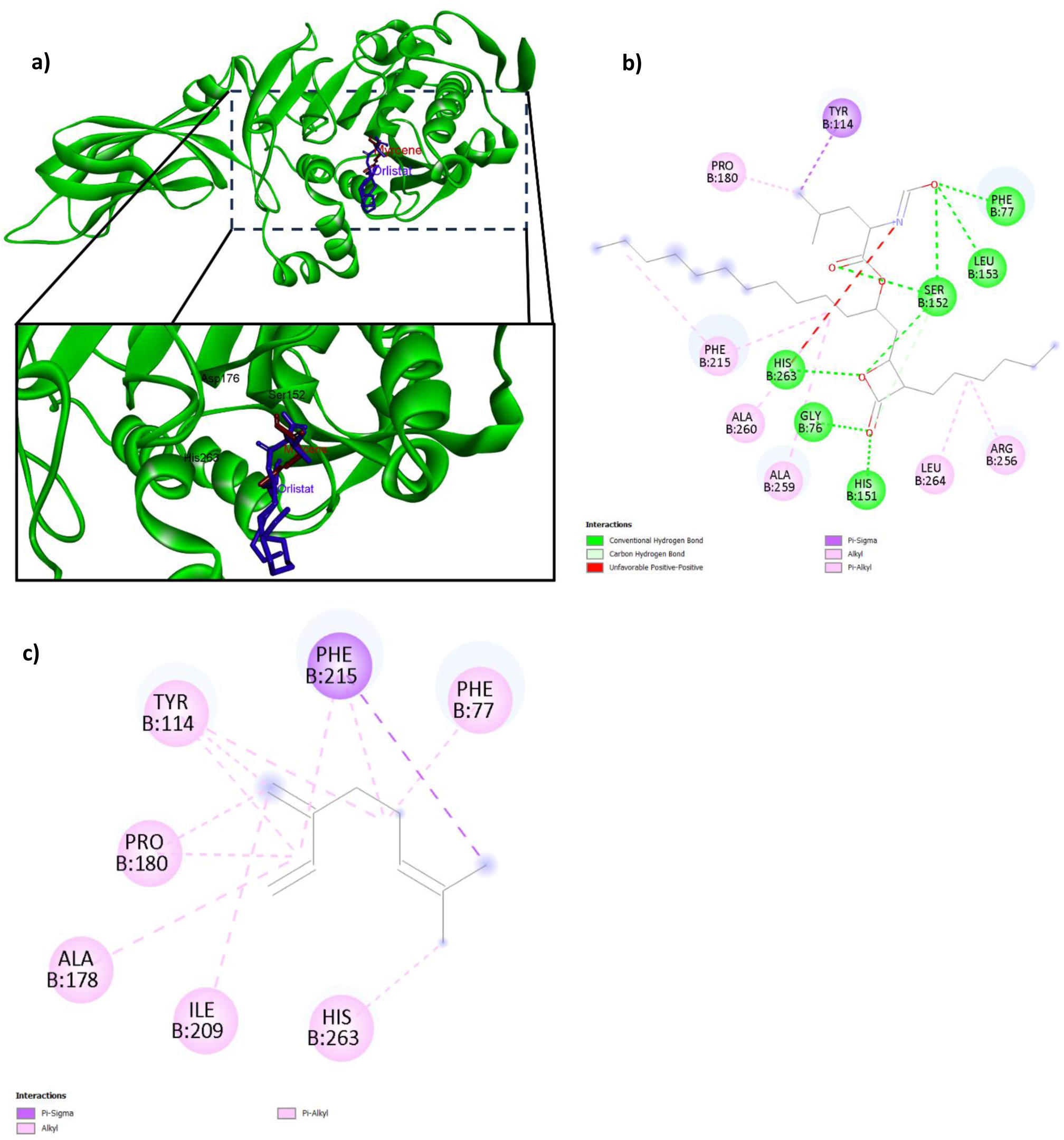
Molecular docking analysis of myrcene (ligand) and orlistat (ligand) on lipase (protein). (a) Binding pocket of myrcene and orlistat on lipase. The catalytic triad (Ser152, Asp176 and His263) are labelled. (b) 2D representation of orlistat (ligand) and lipase (protein) interaction. (c) 2D representation of myrcene (ligand) and lipase (protein) interaction.

### Molecular docking analysis of NF-κB with MCN

Across all ten molecular docking analysis performed with different random seeds, myrcene consistently docked in proximity of the nuclear localization signal motif (Lys301, Arg302, Lys303 and Arg304) (Fig. 8a) of NF-κB P65 (Chain A), exhibiting the highest binding affinity of −5.2 kcal/mol. Myrcene directly interacted with an amino acid residue within the nuclear localization signal motif (Arg302) through either alkyl or π-alkyl interaction (Fig. 8b).

**Fig. 8.**
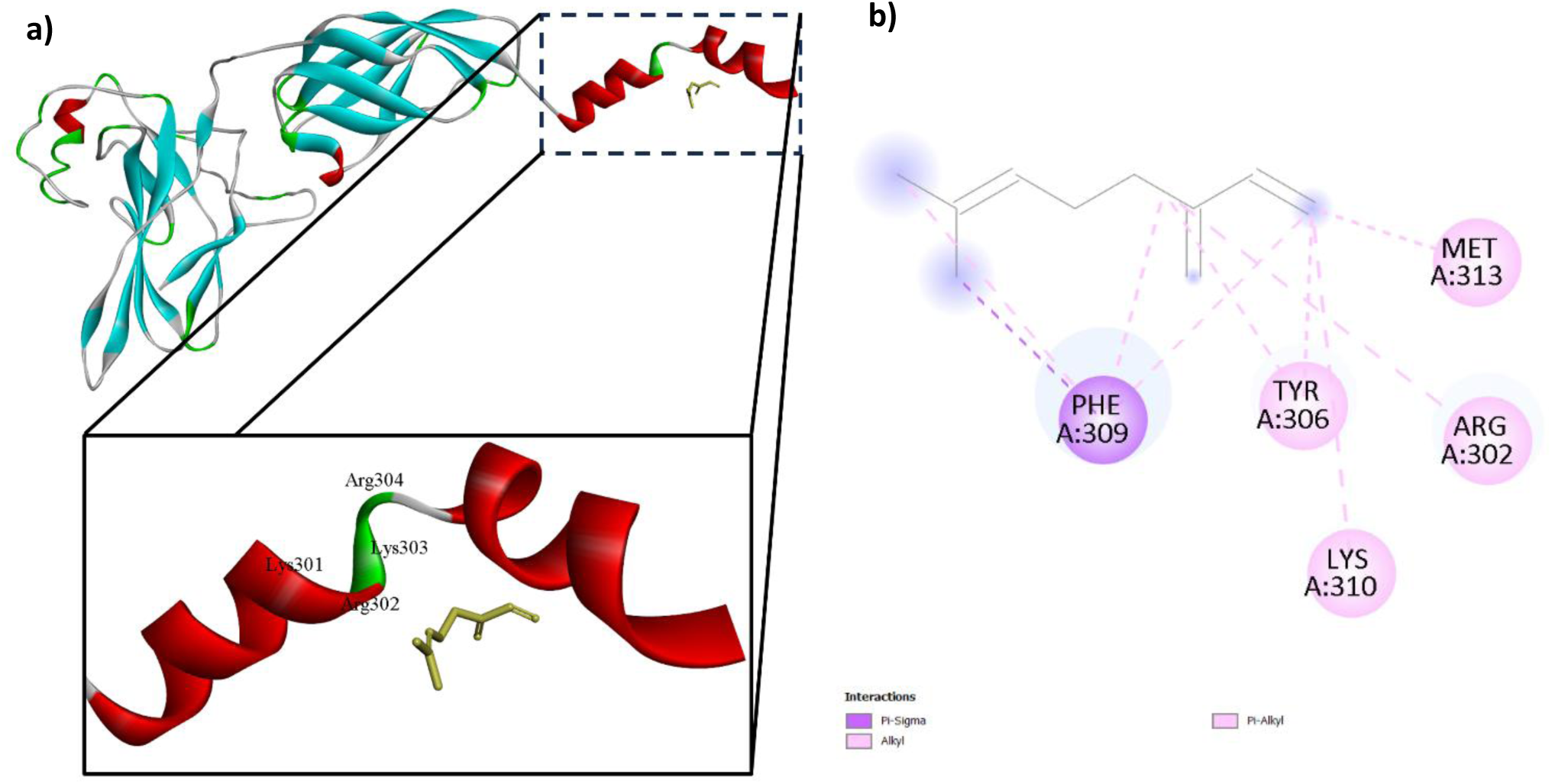
Molecular docking analysis of myrcene (ligand) on NF-κB P65 (protein). (a) Molecular docking of myrcene to the NF-κB P65. The nuclear localization signal motifs (Lys301, Arg302, Lys303, and Arg304) are labelled. (b) 2D representation of myrcene (ligand) and NF-κB P65 (protein) interaction.

## Discussion

The current investigation demonstrated that myrcene (MCN), a naturally occurring monoterpene hydrocarbon, exerts a multifaceted biological role by inhibiting pancreatic lipase and attenuating LPS-induced inflammatory responses in human monocytic THP-1 cells. The dual pharmacological activity of MCN provides an integrated approach toward managing obesity-associated inflammation and metabolic disorders, both of which share overlapping molecular pathways.

Pancreatic lipase plays a central role in the digestion of dietary fat, hydrolyzing triglycerides into free fatty acids and monoglycerides. However, excessive lipase activity results in elevated circulating free fatty acids (FFAs), which contribute to the activation of pro-inflammatory cytokine such as TNF-α in monocytes/macrophages [23]. This lipotoxic response promotes chronic low-grade inflammation, a hallmark of obesity and metabolic syndrome [24]. The present study revealed that MCN inhibited lipase activity in a concentration-dependent manner, corroborating earlier evidence that plant-derived terpenes and flavonoids act as natural lipase inhibitors [25,26].

Docking analysis further confirmed the molecular basis of this inhibition, with MCN forming interactions at the catalytic triad site (His263) of pancreatic lipase, analogous to orlistat’s known binding pattern [22]. Similar observations were made for other terpenoids, such as limonene and pinene, which inhibit pancreatic lipase through hydrophobic and π–π stacking interactions at the catalytic pocket [27]. Thus, MCN may modulate lipid absorption and indirectly reduce inflammatory responses mediated by FFA-induced TLR activation.

Myrcene has long been recognized for its potent anti-inflammatory and antioxidant properties [28]. In the present study, MCN significantly suppressed TNF-α expression and secretion in LPS-stimulated THP-1 cells, reduced TLR2 and MIP-1 gene expression, and inhibited NF-κB p65 nuclear translocation. These findings are consistent with prior work demonstrating that myrcene and other monoterpenes downregulate pro-inflammatory cytokines such as TNF-α, IL-1β, and IL-6 in macrophages, keratinocytes, and chondrocytes [29,30].

The inhibition of NF-κB activation represents a key mechanism in the anti-inflammatory action of MCN. NF-κB is a pivotal transcription factor controlling the expression of numerous inflammatory mediators. Molecular docking analysis revealed that MCN binds near the nuclear localization signal (NLS) region of NF-κB p65 (Arg302), which may hinder the translocation of the NF-κB complex into the nucleus and thereby block inflammatory gene transcription. Similar binding behaviour has been reported for other phytochemicals such as curcumin, carvacrol, and α-pinene, which interfere with the NLS region to suppress NF-κB activation [31,32]. This interaction underscores the potential of MCN to act as a direct modulator of NF-κB signaling at the molecular level.

Computational modeling in this study supports the dual binding affinity of MCN toward both lipase and NF-κB targets. MCN exhibited binding affinities of −5.6 kcal/mol with pancreatic lipase and −5.2 kcal/mol with NF-κB, suggesting that its moderate yet consistent affinity could be sufficient to modulate biological activity, particularly under physiological accumulation or synergistic phytochemical conditions. Previous molecular docking investigations on other monoterpenes (e.g., menthol, thymol, eucalyptol) also reported comparable binding energies with NF-κB and COX-2, which correlate with observed anti-inflammatory outcomes [33–35].

A growing body of literature highlights the intersection between metabolic and inflammatory disorders, where dysregulated lipid metabolism acts as a trigger for chronic inflammation [36]. In this context, natural compounds like myrcene offer therapeutic promise by targeting both processes simultaneously. Studies have demonstrated that terpenoids such as β-caryophyllene and linalool not only inhibit lipase but also suppress NF-κB activation and cytokine production, supporting their dual regulatory potential [37–39]. Similarly, in high-fat diet-induced models, dietary terpenes ameliorated lipid accumulation and systemic inflammation through modulation of AMPK and NF-κB signaling [40]. These observations align closely with the present findings, indicating that MCN might confer comparable metabolic and anti-inflammatory benefits.

The convergence of anti-lipase and anti-inflammatory activities positions MCN as a promising candidate for developing therapeutic interventions against metabolic inflammation, obesity, and associated disorders. Its favorable lipophilicity, natural abundance, and known safety profile enhance its pharmaceutical potential [41]. Nevertheless, challenges such as poor water solubility and limited bioavailability need to be addressed. Encapsulation in nanoemulsions or lipid-based carriers could enhance its therapeutic efficacy, as demonstrated for other terpenes [42]. Furthermore, combinatorial therapy involving MCN and standard anti-inflammatory or lipid-lowering agents may produce synergistic effects with reduced side effects.

## Conclusion

The present study demonstrates that myrcene (MCN), a naturally occurring monoterpene, exhibits dual biological activities by inhibiting pancreatic lipase and suppressing LPS-induced inflammatory responses in human monocytic THP-1 cells. The inhibition of lipase activity by MCN suggests its potential role in regulating lipid metabolism and reducing free fatty acid-induced inflammation. Concurrently, its ability to downregulate TNF-α, TLR2, and MIP-1 expression, as well as inhibit NF-κB p65 nuclear translocation, highlights its strong anti-inflammatory efficacy. Molecular docking studies further supported these findings, revealing that MCN interacts directly with the catalytic triad of lipase and the nuclear localization signal region of NF-κB, indicating a mechanistic basis for its bioactivity. Together, these results suggest that MCN acts as a multi-target modulator that bridges metabolic and inflammatory pathways, offering therapeutic potential against metabolic inflammation, obesity-related disorders, and chronic inflammatory diseases.

## Acknowledgements

Author Smruti extends his appreciation to Late Prof. Sarada D. Tetali for her timely support and valuable suggestions.

## AI declaration statement

The authors used an AI tool (Grammarly AI) to assist in improving the clarity and readability of the manuscript.

## CRediT authorship contribution statement

**Smruti Ranjan Das** contributed to the study design, investigation, and drafted the original manuscript. **Satyabrata Acharya**, **Sachidanand Nayak**, and **Asenath Shishak** contributed to methodology, data visualization, and formal analysis. **Aditya Ankari** contributed to the formal analysis and revision of the manuscript. All authors read and approved the final manuscript.

## Data availability

Data available upon request.

## References

[1] Yoon, J. H., & Baek, S. J. (2005). Molecular targets of dietary polyphenols with anti-inflammatory properties. Yonsei Medical Journal, 46(5), 585–596.

[2] Chen, L., Deng, H., Cui, H., Fang, J., Zuo, Z., Deng, J., … & Zhao, L. (2017). Inflammatory responses and inflammation-associated diseases in organs. Oncotarget, 9(6), 7204.

[3] Kany, S., Vollrath, J. T., & Relja, B. (2019). Cytokines in inflammatory disease. International Journal of Molecular Sciences, 20(23), 1–31.

[4] Negus, R. P. M. (1996). The chemokines: Cytokines that direct leukocyte migration.. Journal of the Royal Society of Medicine, 89(6), 312–314.

[5] Kumar, A. N., Das, S. R., Kumar, J. K., Srinivas, K. V. N. S., & Tetali, S. D. (2025). Design and development of an isatin-1,2,3-triazole hybrid analogue as a potent anti-inflammatory agent with enhanced efficacy and gene expression modulation. RSC Advances, 15(3), 2023–2033.

[6] Libby, P. (2012). Inflammation in atherosclerosis. *Arteriosclerosis*, Thrombosis, and Vascular Biology, 32(9), 2045–2051.

[7] Reddi, K. K., & Tetali, S. D. (2019). Dry leaf extracts of Tinospora cordifolia (Willd.) Miers attenuate oxidative stress and inflammatory condition in human monocytic (THP-1) cells. Phytomedicine, 61, 152831.

[8] Gesell, A., Chávez, M. L. D., Kramell, R., Piotrowski, M., Macheroux, P., & Kutchan, T. M. (2011). Heterologous expression of two FAD-dependent oxidases with (S)-tetrahydroprotoberberine oxidase activity from *Argemone mexicana* and *Berberis wilsoniae* in insect cells. Planta, 233(6), 1185–1197.

[9] Mukaila, Y. O., Pfukwa, T. M., & Fawole, O. A. (2025). Essential oils from South African indigenous plants: Extraction techniques, phytochemistry, biological activities and applications. South African Journal of Botany, 180, 774–794.

[10] Abdel-Rahman, R. F., Soliman, G. A., Saeedan, A. S., Ogaly, H. A., Abd-Elsalam, R. M., Alqasoumi, S. I., & Abdel-Kader, M. S. (2019). Molecular and biochemical monitoring of the possible herb-drug interaction between *Momordica charantia* extract and glibenclamide in diabetic rats. Saudi Pharmaceutical Journal, 27(6), 803–816.

[11] Dragoș, D., Enache, I. I., & Manea, M. M. (2025). Oxidative stress and nutritional antioxidants in renal diseases: A Narrative Review. Antioxidants, 14(7), 1–33.

[12] Ashruf, O. S., & Ansari, M. Y. (2023). Natural compounds: potential therapeutics for the inhibition of cartilage matrix degradation in osteoarthritis. Life, 13(1), 1–20.

[13] Chanda, J., Mukherjee, P. K., Biswas, R., Malakar, D., & Pillai, M. (2019). Study of pancreatic lipase inhibition kinetics and LC–QTOF–MS-based identification of bioactive constituents of *Momordica charantia* fruits. Biomedical Chromatography, 33(4), 1–9.

14. Gonçalves, R., Mateus, N., & de Freitas, V. (2010). Study of the interaction of pancreatic lipase with procyanidins by optical and enzymatic methods. Journal of Agricultural and Food Chemistry, 58(22), 11901–11906.

[15] Choudhury, S. S., Bashyam, L., Manthapuram, N., Bitla, P., Kollipara, P., & Tetali, S. D. (2014). *Ocimum sanctum* leaf extracts attenuate human monocytic (THP-1) cell activation. Journal of Ethnopharmacology, 154(1), 148–155.

[16] Reddi, K. K., Li, H., Li, W., & Tetali, S. D. (2021). Berberine, a phytoalkaloid, inhibits inflammatory response induced by lps through nf-kappaβ pathway: Possible involvement of the ikkα. Molecules, 26(16).

[17] Kokkiripati, P. K., Kamsala, R. V., Bashyam, L., Manthapuram, N., Bitla, P., Peddada, V., Raghavendra, A. S., & Tetali, S. D. (2013). Stem-bark of *Terminalia arjuna* attenuates human monocytic (THP-1) and aortic endothelial cell activation. Journal of Ethnopharmacology, 146(2).

[18] Dong, W., Wang, X., Bi, S., Pan, Z., Liu, S., Yu, H., Lu, H., Lin, X. I., Wang, X., Ma, T., & Zhang, W. (2014). Inhibitory effects of resveratrol on foam cell formation are mediated through monocyte chemotactic protein-1 and lipid metabolism-related proteins. International Journal of Molecular Medicine, 33(5), 1161–1168.

[19] Morris, G. M., Huey, R., Lindstrom, W., Sanner, M. F., Belew, R. K., Goodsell, D. S., & Olson, A. J. (2009). AutoDock4 and AutoDockTools4: Automated docking with selective receptor flexibility. Journal of Computational Chemistry, 30(16), 2785–2791.

[20] O’Boyle, N. M., Banck, M., James, C. A., Morley, C., Vandermeersch, T., & Hutchison, G. R. (2011). Open Babel: An open chemical toolbox. Journal of Cheminformatics, 3(1), 33.

[21] Trott, O., & Olson, A. J. (2010). AutoDock Vina: improving the speed and accuracy of docking with a new scoring function, efficient optimization, and multithreading. Journal of Computational Chemistry, 31(2), 455–461.

[22] Gholami, A., Minai-Tehrani, D., & Eriksson, L. A. (2025). Combining kinetics and in silico approaches to evaluate bromhexine as an anti-pancreatic lipase agent for obesity management. Scientific Reports, 15(1), 1–12.

[23] Kojta, I., Chacińska, M., & Błachnio-Zabielska, A. (2020). Obesity, bioactive lipids, and adipose tissue inflammation in insulin resistance. Nutrients, 12(5), 1305.

[24] Lafontan, M., & Langin, D. (2009). Lipolysis and lipid mobilization in human adipose tissue. Progress in Lipid Research, 48(5), 275–297.

[25] Birari, R. B., & Bhutani, K. K. (2007). Pancreatic lipase inhibitors from natural sources: unexplored potential. Drug Discovery Today, 12(19–20), 879–889.

[26] He, L., Su, Z., & Wang, S. (2024). The anti-obesity effects of polyphenols: a comprehensive review of molecular mechanisms and signal pathways in regulating adipocytes. Frontiers in Nutrition, 11(October), 1–20.

[27] Wang, P., Song, X., & Liang, Q. (2025). Molecular docking studies and in vitro activity of pancreatic lipase inhibitors from yak milk cheese. International Journal of Molecular Sciences, 26(2), 756.

[28] De Cássia Da Silveira E Sá, R., Andrade, L. N., & De Sousa, D. P. (2013). A review on anti-inflammatory activity of monoterpenes. Molecules, 18(1), 1227–1254.

[29] Almarzooqi, S., Venkataraman, B., Raj, V., Alkuwaiti, S. A. A., Das, K. M., Collin, P. D., Adrian, T. E., & Subramanya, S. B. (2022). β-Myrcene mitigates colon inflammation by inhibiting MAP kinase and NF-κB signaling pathways. Molecules, 27(24).

[30] Quintans, J. S., Shanmugam, S., Heimfarth, L., Araújo, A. A. S., Almeida, J. R. D. S., Picot, L., & Quintans-Júnior, L. J. (2019). Monoterpenes modulating cytokines-A review. Food and Chemical Toxicology, 123, 233–257.

[31] Biswas, D., Somkuwar, B. G., Borah, J. C., Varadwaj, P. K., Gupta, S., Khan, Z. A., Mondal, G., Chattoraj, A., & Deb, L. (2023). Phytochemical mediated modulation of COX-3 and NFκB for the management and treatment of arthritis. Scientific Reports, 13(1), 1–14.

[32] Katsori, A. M., Palagani, A., Bougarne, N., Hadjipavlou-Litina, D., Haegeman, G., & Berghe, W. Vanden. (2015). Inhibition of the NF-κB signaling pathway by a novel heterocyclic curcumin analogue. Molecules, 20(1), 863–878.

[33] Islam, M. T., Bappi, M. H., Bhuia, M. S., Ansari, S. A., Ansari, I. A., Shill, M. C., … & El-Nashar, H. A. (2024). Anti-inflammatory effects of thymol: an emphasis on the molecular interactions through in vivo approach and molecular dynamic simulations. Frontiers in Chemistry, 12, 1376783.

[34] Mohamed, M. E., Aldhubiab, B., & Younis, N. S. (2025). Eucalyptol Promotes Heart Restoration in Rats Following Myocardial Infarction Through TLR/NF-κB. Pharmacognosy Magazine, 09731296251361321.

[35] Wang, X., Lai, J., Xu, F., & Liu, M. (2025). Network pharmacology and molecular docking: exploring the mechanism of peppermint in mastitis prevention and treatment in dairy cows. Veterinary Sciences, 12(2), 1–19.

[36] Hotamisligil, G. S. (2017). Inflammation, metaflammation and immunometabolic disorders. Nature, 542(7640), 177–185.

[37] Li, Y., Lv, O., Zhou, F., Li, Q., Wu, Z., & Zheng, Y. (2015). Linalool inhibits LPS-induced inflammation in BV2 microglia cells by activating Nrf2. Neurochemical Research, 40(7), 1520–1525.

[38] Rodrígue z-Mejía, U. U., Viveros-Paredes, J. M., Zepeda-Morales, A. S. M., Carrera-Quintanar, L., Zepeda-Nuño, J. S., Velázquez-Juárez, G., Delgado-Rizo, V., García-Iglesias, T., Camacho-Padilla, L. G., Varela-Navarro, E., Anguiano-Sevilla, L. A., Franco-Torres, E. M., & López-Roa, R. I. (2022). β-Caryophyllene: A therapeutic alternative for intestinal barrier dysfunction caused by obesity. Molecules, 27(19).

[39] Wei, Z., Chen, G., Hu, T., Mo, X., Hou, X., Cao, K., Wang, L., Pan, Z., Wu, Q., Li, X., Ye, F., Zouboulis, C. C., & Ju, Q. (2021). Resveratrol ameliorates lipid accumulation and inflammation in human SZ95 sebocytes via the AMPK signaling pathways in vitro. Journal of Dermatological Science, 103(3), 156–166.

[40] Lindholm, C. R., Ertel, R. L., Bauwens, J. D., Schmuck, E. G., Mulligan, J. D., & Saupe, K. W. (2013). A high-fat diet decreases AMPK activity in multiple tissues in the absence of hyperglycemia or systemic inflammation in rats. Journal of Physiology and Biochemistry, 69(2), 165–175.

[41] Surendran, S., Qassadi, F., Surendran, G., Lilley, D., & Heinrich, M. (2021). Myrcene—what are the potential health benefits of this flavouring and aroma agent?. Frontiers in Nutrition, 8, 699666.

[42] Nikolic, I., Mitsou, E., Pantelic, I., Randjelovic, D., Markovic, B., Papadimitriou, V., Xenakis, A., Lunter, D. J., Zugic, A., & Savic, S. (2020). Microstructure and biopharmaceutical performances of curcumin-loaded low-energy nanoemulsions containing eucalyptol and pinene: Terpenes’ role overcome penetration enhancement effect? European Journal of Pharmaceutical Sciences, 142, 105135.

