## Supplementary figures for "Integrative analysis of Myrcene’s lipase inhibition and anti-inflammatory mechanisms in human monocytic cells"

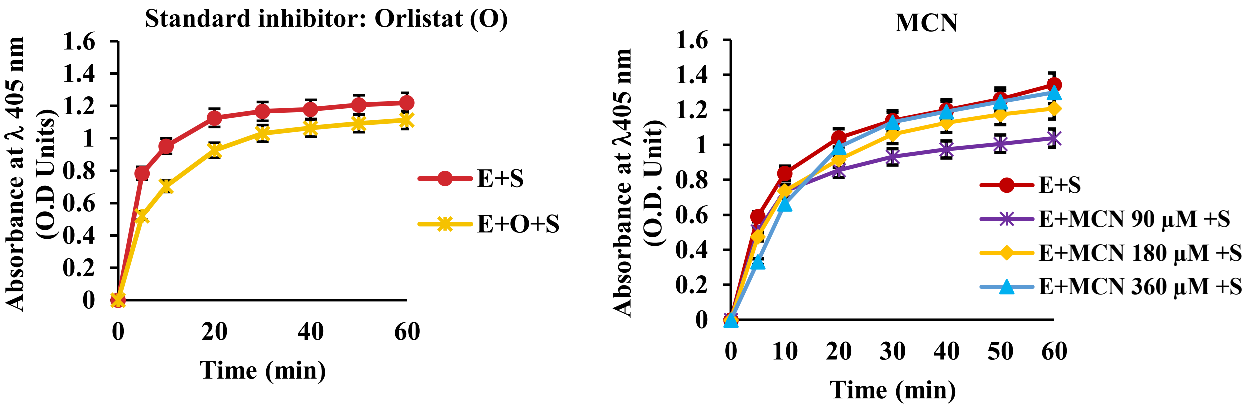


**Fig.1.** Inhibitory effect of (a) orlistat and (b) selected metabolite myrcene (MCN) on pancreatic lipase enzyme. Data are presented as mean ± standard deviation (SD) from three independent experiments (n=3); statistical significance was set at P< 0.001.

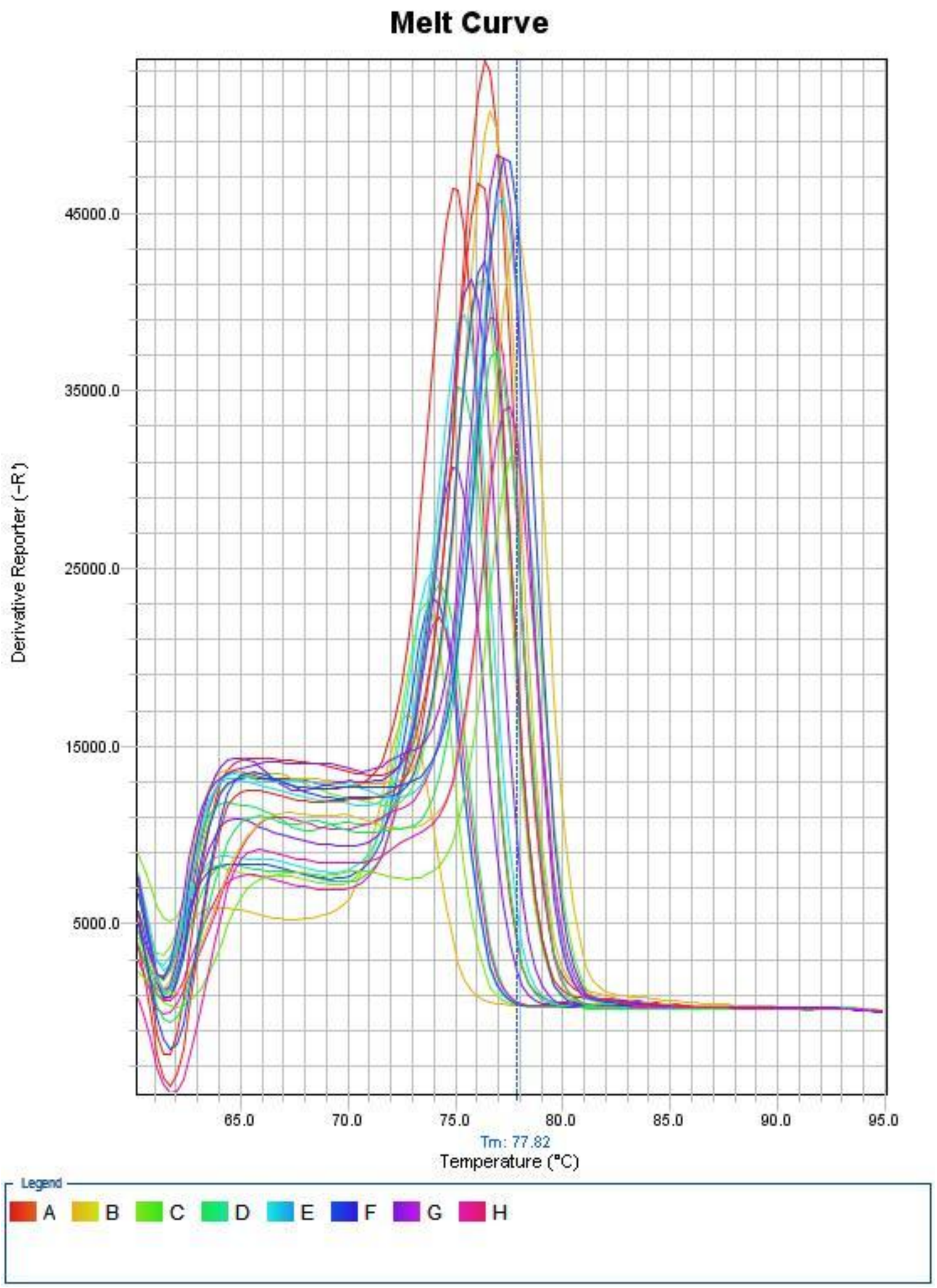


**Fig.2.** Melt curve analysis of the GAPDH RT-PCR product in THP-1 cells treated with MCN and stimulated with LPS.

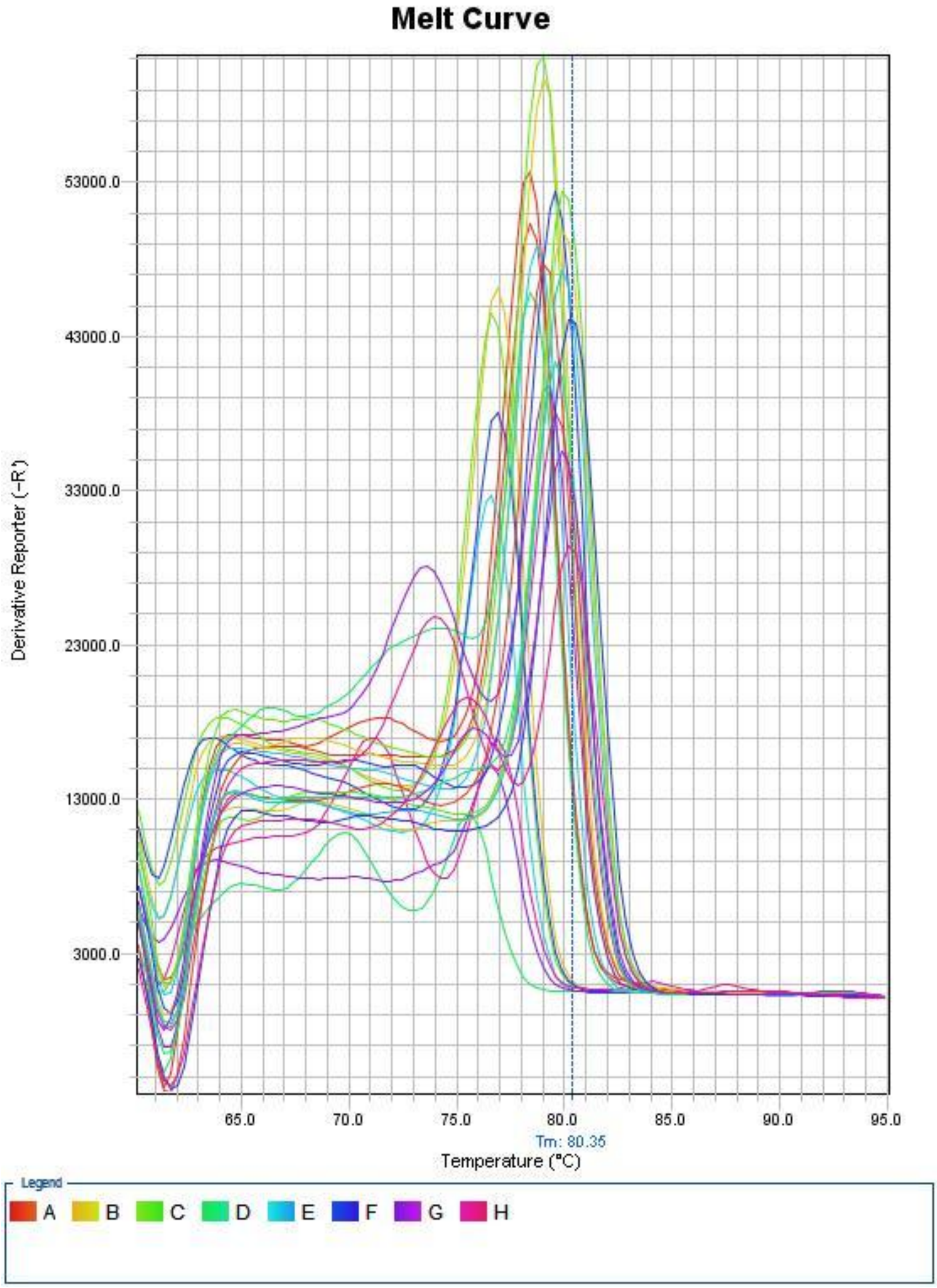


**Fig.3.** Melt curve analysis of the TNF-α RT-PCR product in THP-1 cells treated with MCN and stimulated with LPS.

a) b)

c) d)

e) f)

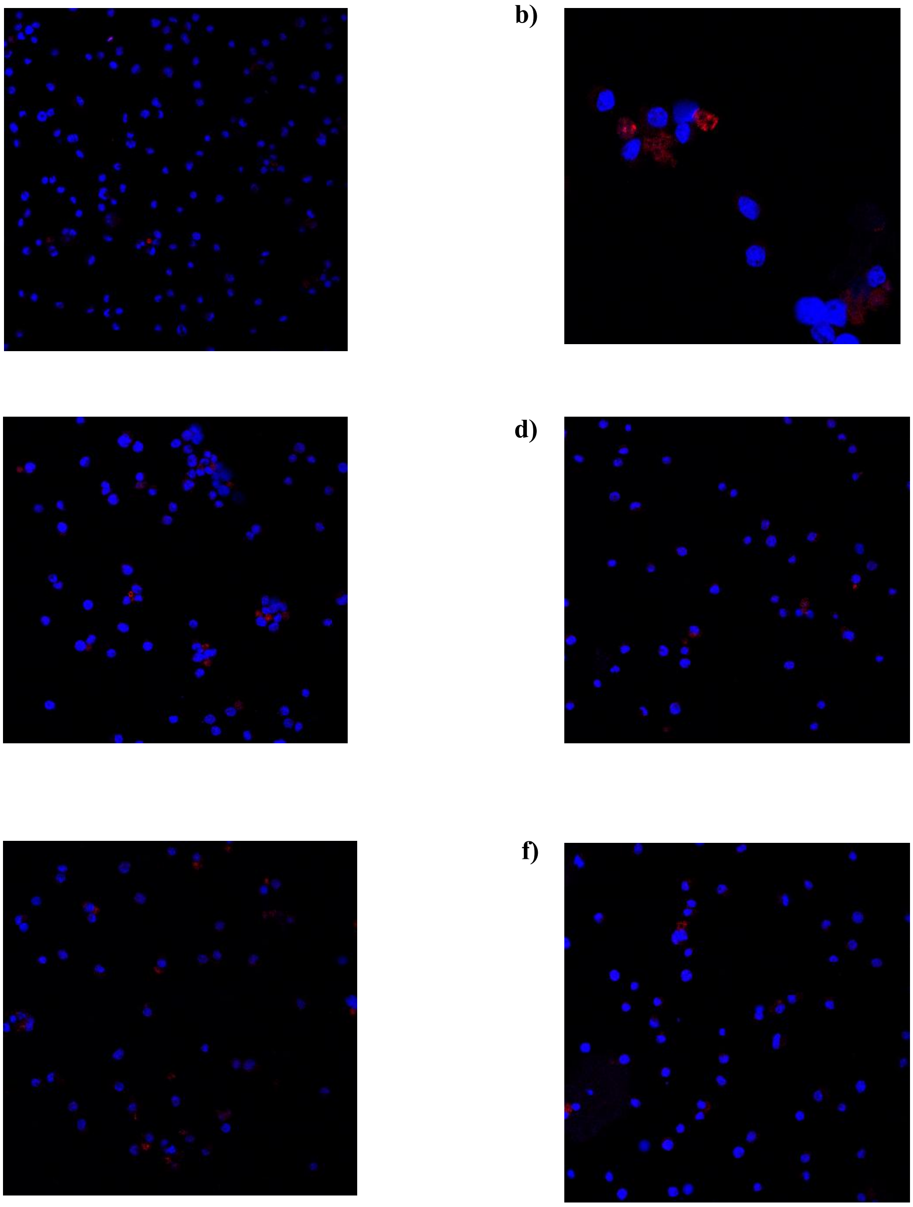

**Fig.4.** Effect of MCN on LPS-induced NF-κB p65 subunit translocation. THP-1 cells were stimulated with LPS (0.5 μg/mL), and the subcellular localization of the p65 subunit was assessed using immunofluorescence. NF-κB p65 was detected with an Alexa Fluor 594-conjugated secondary antibody, and nuclei were counterstained with DAPI. Images were acquired using a Leica confocal microscope. Panels represent: (a) Cells, (b) Cells + LPS, (c) Cells + MCN (36 μM) + LPS, (d) Cells + MCN (72 μM) + LPS, (e) Cells + MCN (108 μM) + LPS, and (f) Cells + MCN (144 μM).

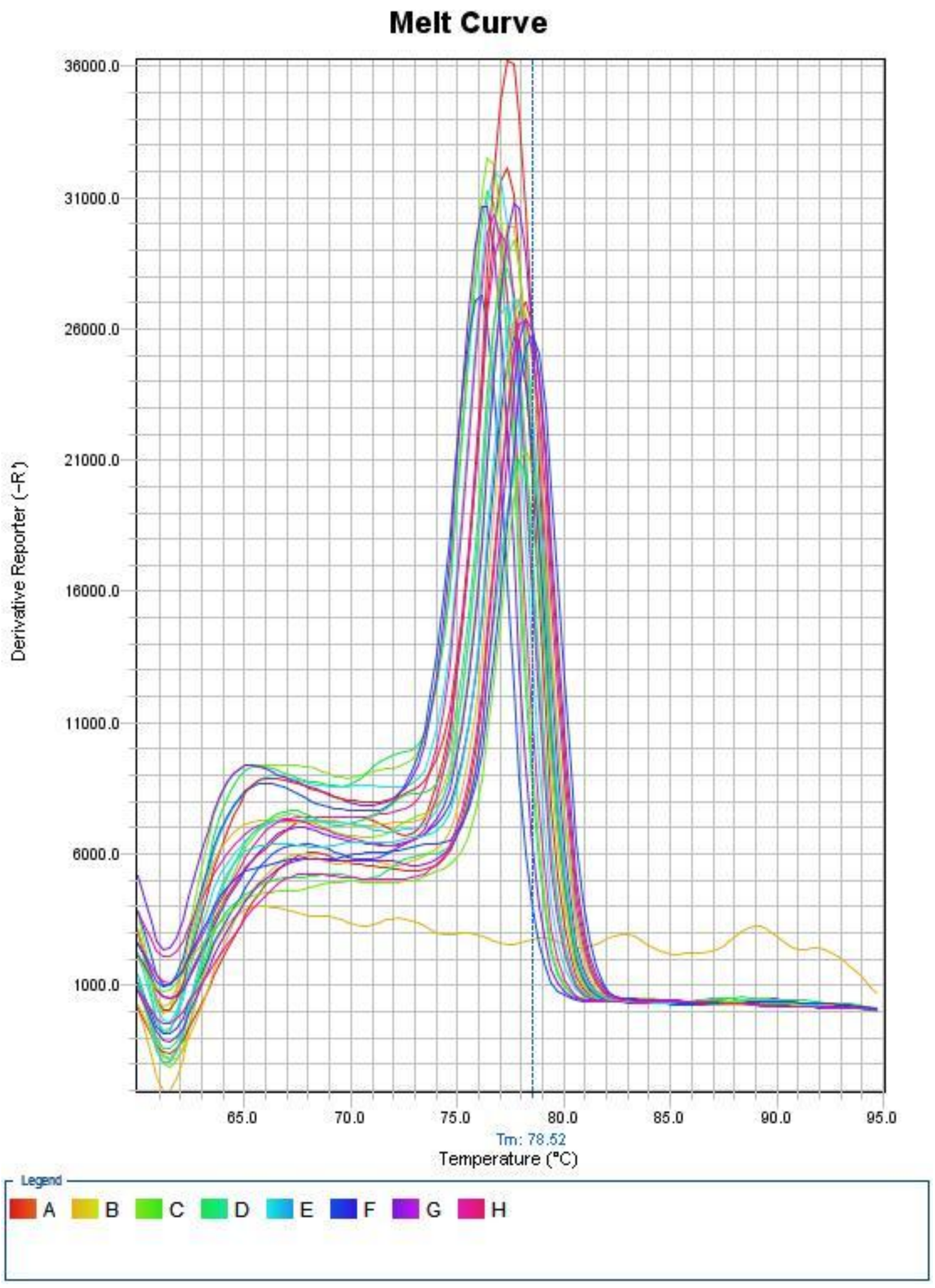


**Fig.5.** Melt curve analysis of the GAPDH RT-PCR product in THP-1 cells treated with MCN and stimulated with PMA.

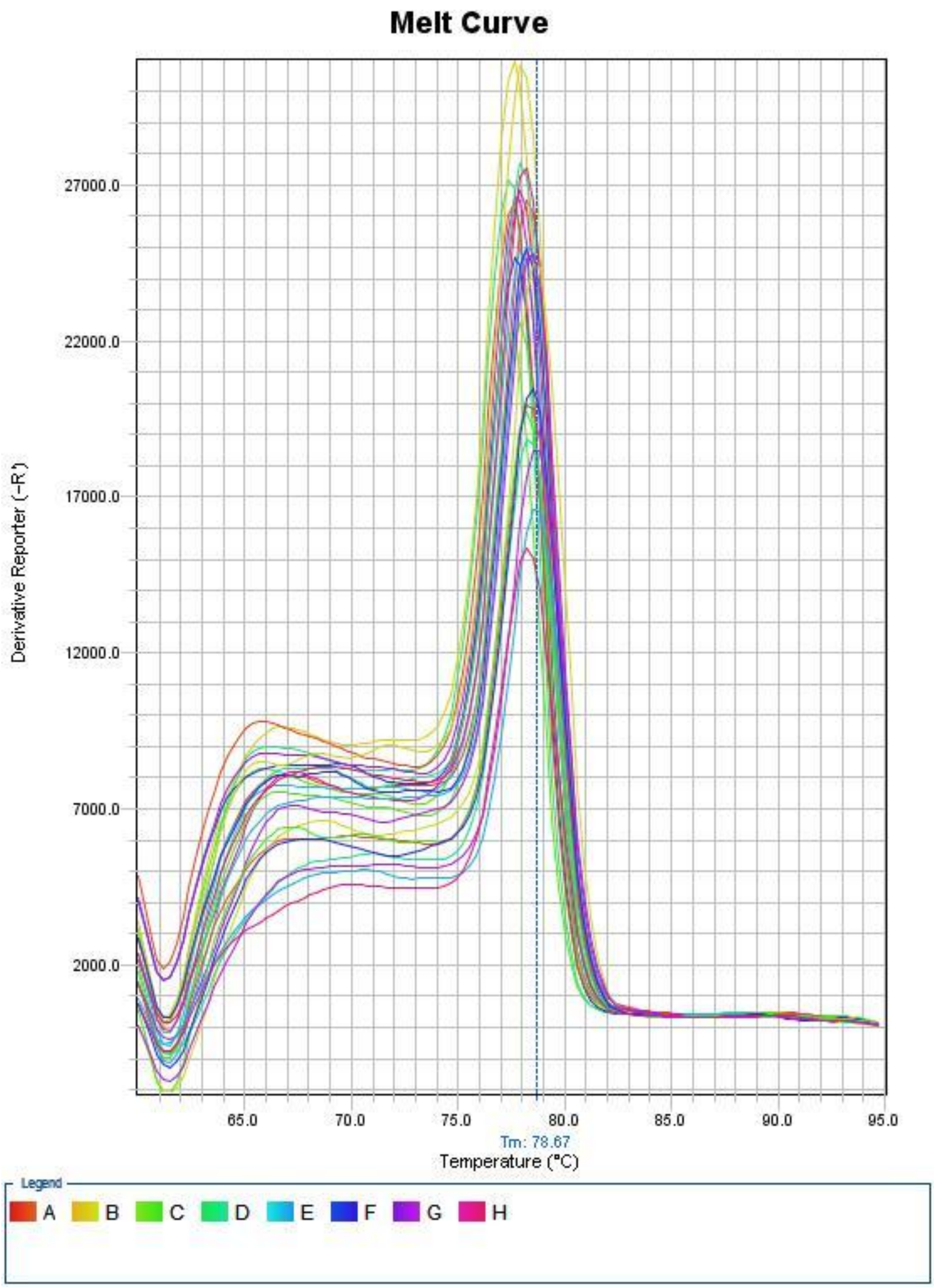


**Fig.6.** Melt curve analysis of the MIP-1 RT-PCR product in THP-1 cells treated with MCN and stimulated with PMA.

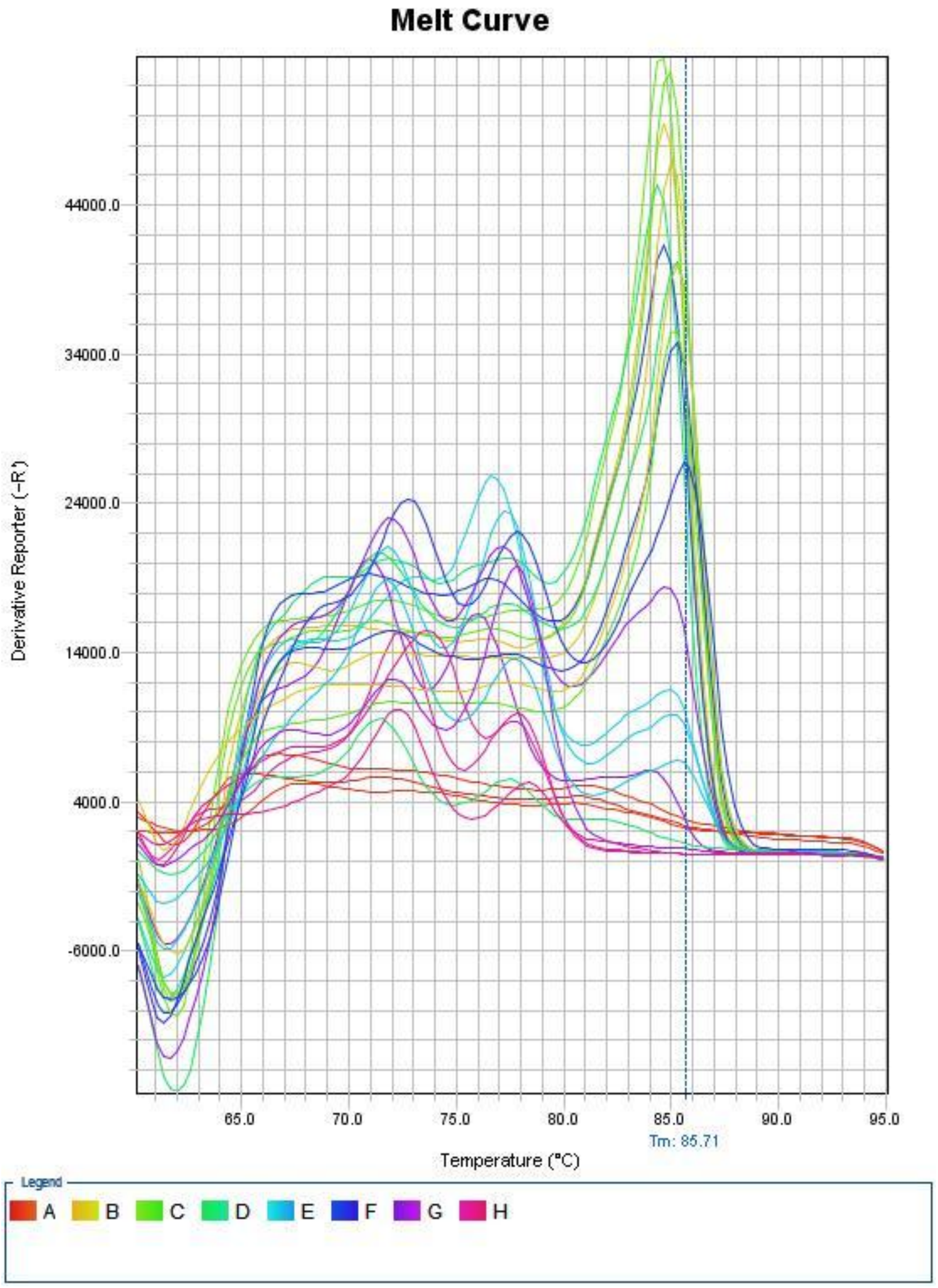


**Fig.7.** Melt curve analysis of the TLR2 RT-PCR product in THP-1 cells treated with MCN and stimulated with PMA.
